# QuASAR: Quality Assessment of Spatial Arrangement Reproducibility in Hi-C Data

**DOI:** 10.1101/204438

**Authors:** Michael EG Sauria, James Taylor

## Abstract

Hi-C has revolutionized global interrogation of chromosome conformation, however there are few tools to assess the reliability of individual experiments. Here we present a new approach, QuASAR, for measuring quality within and between Hi-C samples. We show that QuASAR can detect even tiny fractions of noise and estimate both return on additional sequencing and quality upper bounds. We also demonstrate QuASAR's utility in measuring replicate agreement across feature resolutions. Finally, QuASAR can estimate resolution limits based on both internal and replicate quality scores. QuASAR provides an objective means of Hi-C sample comparison while providing context and limits to these measures.

## Background

Chromosome Conformation Capture assays, particularly Hi-C, are becoming widely used, and have enabled a much better understanding of chromatin spatial interactions and architecture. What has not kept pace is the ability to assess the reliability of these data. One dimensional genomic annotation assays, such as ChIP-seq, have been the focus of quality control efforts for years[1–6]. These methods take a variety of approaches for assessing sample quality including external control samples, orthogonal datasets, replicate comparison, and mapping statistics. This allows direct comparison between similar datasets and a degree of confidence in the biological conclusions associated with the results of analyzing these datasets. The field of chromatin topology would benefit from a similar set of quality assessment resources.

Three areas need to be addressed when considering Hi-C data quality. Two of these, individual sample quality and replicate agreement, are also of great importance in other genomic annotation data types. The final area, limits of data resolution, is a consideration in other types of genomic data but has a special importance in Hi-C data because of the two dimensional nature of the data compared to more traditional one dimensional annotation data. For example, the amount of sequencing required to achieve saturation in a mammalian ChIP-seq experiment is about 40 million reads [7]. While not trivial, this depth of sequencing is often achieved. It is currently unknown what the saturation point of a Hi-C experiment is, but given that most Hi-C samples rarely have more than a couple hundred million reads prior to mapping and filtering, it is unlikely that they approach the saturation point for 2D dataset. Thus it is important to be able to determine reasonable limits for analysis resolution.

A variety of approaches have been use to infer Hi-C sample quality, primarily focused on statistics derived from sequencing quality, read alignment and the position of read ends relative to restriction fragments [8–14]. Sequencing quality and alignment quality give some information about sample integrity and possible contamination. Rao et al. [11] proposed a complexity statistic to determine which Hi-C libraries were worth sequencing. This statistic was based on the percent of unique reads in the sequenced sample. Other groups have pre-screened samples using fragment size profiles as an indication of Hi-C library quality prior to sequencing [12, 13]. While these approaches can provide information about the utility of sample processing, they fail to provide information about how well the Hi-C library captures chromatin conformations reflective of the ground truth. Statistics derived from aligned reads such as PCR duplication rates, self-ligation of restriction fragments, or the percentage reads with inserts of expected size can also be useful in determining the percentage reads in a Hi-C library that have been correctly processed, and it is crucial to perform filtering based on these features. They do not, however, provide information about the actual conformational quality of reads passing these filters, only that the reads conform to a set of expected characteristics based on the Hi-C protocol. For example, a negative control sample lacking the cross-linking step would produce a Hi-C library with the a similar percentage of reads passed to downstream analysis as a normally processed Hi-C sample. The one statistic that has been put forth to directly address sample quality is the ratio of intra- to inter-chromosomal reads [15]. If fragments are randomly ligated, rather than based on proximity, the number of inter-chromosomal interactions would increase, lowering this ratio. The shortcoming of this measure is that it does not necessarily provide information about the quality of intra-chromosomal reads, nor does it account for possible biologically relevant drivers of increased inter-chromosomal interactions such as chromatin decondensation or release from lamina associated domains.

The second important consideration in Hi-C data quality is reproducibility between samples. A primary means of establishing the similarity between two Hi-C samples has been measuring correlation, either Spearman or Pearson, between heatmaps at a chosen resolution [11, 15–19]. The largest drawback to this approach is that the strongest driver of Hi-C signal is genomic distance between loci, regardless of biological interactions [20]. This means that all Hi-C datasets have the same strong underlying distance-based signal decay driving correlation and differences between samples are small by comparison and harder to detect. Two methods have been devised specifically to address this problem, HiCRep [21] and HiC-Spector [22]. The authors of HiCRep demonstrate that heatmap correlation measures are insufficient at consistently distinguishing unrelated samples from each other compared to biological replicates. This indicates that correlation-based assessments of sample similarity are unreliable. Both of these approaches provide a single summary statistic for the reproducibility between samples, although HiCRep also provides feedback about saturation of the reproducibility measure as a function of sequencing depth.

The final consideration for Hi-C quality assessment is determining what levels of resolution are appropriate for a dataset given its sequencing depth, quality, and reproducibility. Rao et al. [11] proposed a measure of resolution based on minimum number of contacts for some percentage of bins produced by partitioning the genome at a given resolution. This intuitively makes sense from the standpoint of having sufficient data density to resolve features at a given resolution. However, consider a pair of samples with equal numbers of reads, one with a large proportion of random ligation: assuming similar marginal distributions of reads, both samples would have identical resolution limit statistics but different sample qualities at the same resolution.

Here we present a new approach for measuring Hi-C data quality, both within and between samples, using a technique called Quality Assessment of Spatial Arrangement Reproducibility (QuASAR). Using a combination of matrix transformations and sub-sampling, we show that QuASAR not only provides information about a sample’s quality and agreement between replicates, but also estimates for return on additional sequencing, absolute quality limits, and an estimate of the maximum reliable resolution that can be used for a Hi-C sample. Together, these results show that QuASAR can facilitate optimized choices in Hi-C data production as well as informed data comparisons and analysis parameter selection.

## Results

### Spatial consistency concept and matrix transformation strategy

To quantify Hi-C quality, we consider the consistency of inferred spatial arrangement of the Hi-C intra-chromosomal (*cis*) data. Initially, the genome is partitioned into uniform-sized bins at a chosen resolution. For bins that occur close together in space as determined by their read count, there should be a high correlation between their sets of *cis*interactions (Figure 1A). Conversely, bins occurring further apart should show little or no correlation across interactions. Thus, for any given pair of bins, we can identify disagreement between the direct and inferred measure of their interaction. For each sample, we produced a “QuASAR-transformed matrix” by finding the element-wise product of the read count matrix and the local correlation matrix as calculated from a distance-corrected enrichment matrix (Figure 1B). Transformed matrices are calculated across multiple resolutions to examine consistency of different features and scales. In order to target features appropriate to a given resolution, we limit the maximum interaction genomic distance used for analysis as a function of the bin size. This includes in the calculation of correlations, thus the term “local correlation”. The resulting transformed matrices can then be used to calculate individual sample quality scores and replication scores for pairs of samples.

**Figure 1:**
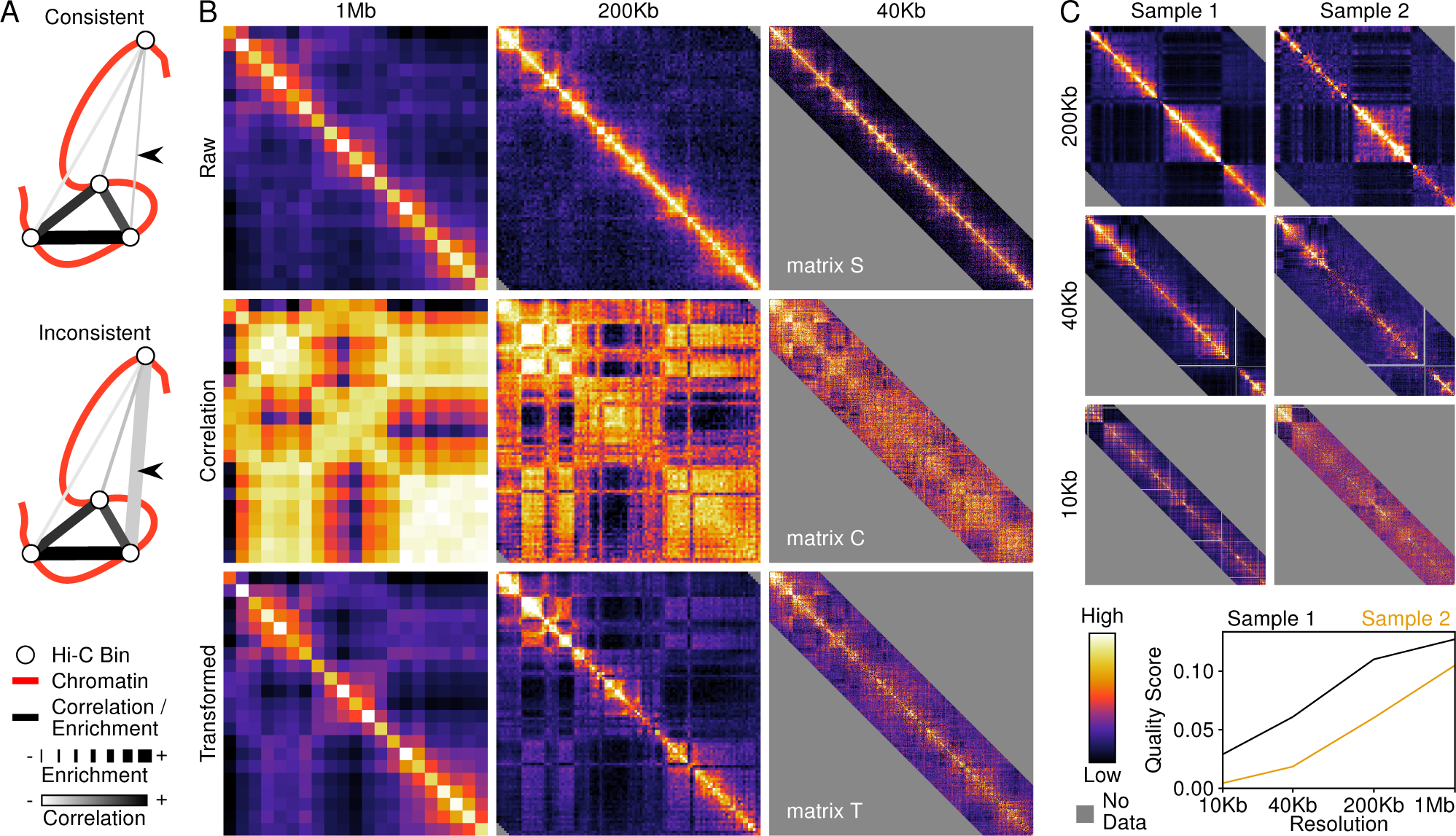
Using spatial consistency between local and regional signals to identify Hi-C sample quality. A) Sample reads are partitioned into bins and pairwise assessment of interaction strength and bin correlation are compared to identify inconsistencies (black arrows). B) Multiple resolutions are considered by QuASAR using transformed matrices derived from local correlation matrices weighted by interaction enrichment. C) Two samples derived from mouse ES cells of differing quality. Lower quality manifests as random points indicating non-specific ligation.

The primary drivers of low quality are random ligation products and missing expected interaction fragments. Within the QuASAR transformed matrix, these types of noise appear as individual entries showing deviation from their surroundings (Figure 1C), higher than local background in the case of random ligation and lower for missing reads. A third factor that may impact the quality score is population heterogeneity, which will manifest as a compression of the dynamic range of signal and less differentiation between the correlation and transformed matrices.

In order to determine an individual sample’s quality, we find the mean of the correlation matrix © weighted by the read count matrix (*R*) minus the unweighted mean (Figure 1B):

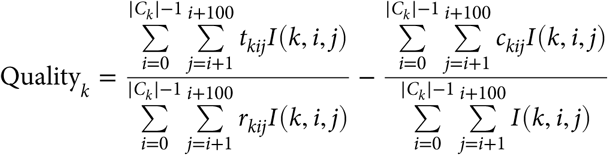

where for bins *i* and *j* on chromosome *k*, *c_kij_* is the correlation from matrix *C_k_*, *r_kij_* is the raw enrichment value from matrix *R_k_*, and *t_kij_* is the transformed value from the QuASAR transformed matrix *T_k_*. *I* is an indicator function taking on a value of one or zero for valid and invalid correlations, respectively.

Replicate scores were determined by finding the correlation of valid values from the transformed matrices (*T_k_*) for chromosome *k* between two samples:

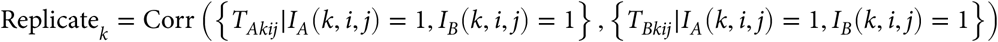

where *I_A_* and *I_B_* are the indicator functions for samples *A* and *B*, respectively. In order to find sample-wide scores, summations and correlations were calculated across all chromosome matrices simultaneously.

### QuASAR evaluation strategy

In order to assess the effectiveness of QuASAR, we used three separate testing approaches. First, we generated noise models for different genome/restriction enzyme combination and created datasets with varying percentages of reads drawn from these models. Second, we used combinations of reads from two separate datasets, either biological replicates in order to generate pseudo-replicates or unrelated samples to create heterogeneous population samples. Finally, we used sub-sampling to investigate the effects of numbers of reads. This was done using *cis*reads that had already passed initial mapping and circularization filters. Because this does not, strictly speaking, qualify as sequencing depth, we refer to the number of valid *cis*reads as “coverage”.

To ensure our results were robust, we tested 96 samples across three species, Mouse, Human, and Drosophila melanogaster (Table S1). Samples ranged in coverage from less than 1 million to 185 million reads. All samples were paired biological replicates and were generated from a diverse set of tissues and cell lines, and originated from numerous laboratories.

### QuASAR Quality results

QuASAR quality scoring effectiveness was tested using injection of simulated random ligation noise, at levels of 0.1% to 75% relative to read coverage. All samples showed a monotonically decreasing relationship between quality signal and percent noise, with the exception of eight instances (Figure 2A). Exceptions to this relationship all occurred at the lowest level of resolution and the majority occurred at under 0.5% noise, with all of them occurring at 5% noise or less. The highest deviating value as a percentage of the raw sample score, was 100.02273%. Because all of these exceptions occurred at low resolution, which is less responsive to coverage and noise effects, and because increases in quality score were minimal, it is likely that these represent stochastic noise. At all but the lowest resolution, as little as 0.1% noise was detectable in every sample using the QuASAR quality score. These results suggest that this metric is sensitive to even small changes in the amounts of random ligation present in Hi-C samples.

**Figure 2:**
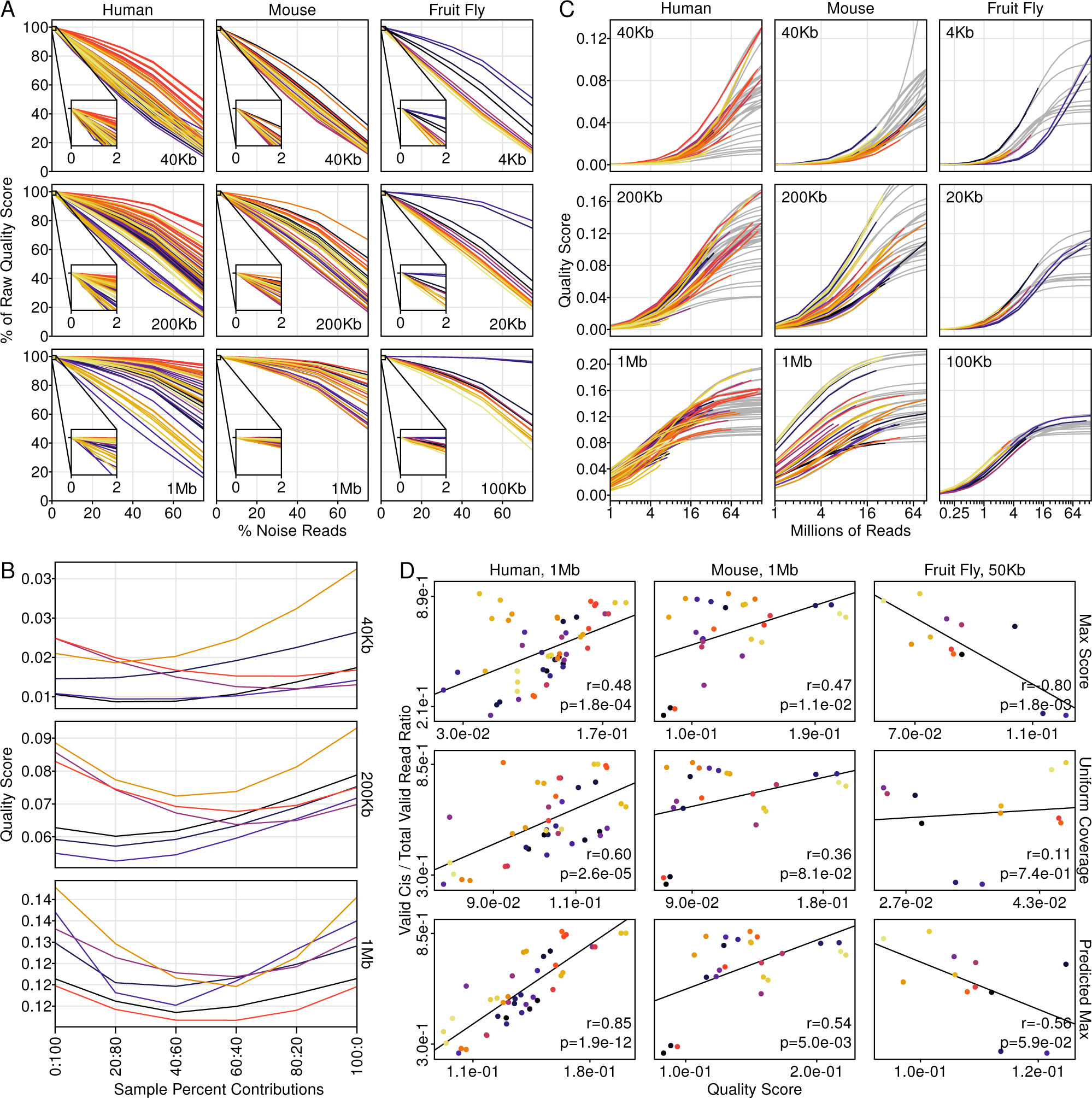
QuASAR quality scoring is sensitive to noise and coverage differences. A) Samples show an almost entirely negative relationship between injected noise and quality score. All scores are reported as a percentage of their unaltered sample score. The insets show a detailed view of values for noise levels between 0 and 2%. All samples within a species are colored consistently across resolutions. B) QuASAR quality scores for mixed samples with varying levels of contribution from each sample pair. All samples have 20 million *cis*reads. C) All quality scores show a monotonically increasing relationship to coverage. Gray lines indicate logistic curves fit to each sample. All samples within a species are colored consistently across resolutions. D) The relationship between quality scores and the percentage intra-chromosomal reads (*cis*) out of all valid reads is depicted. Quality scores were assessed at three different points: the score at maximum sample coverage (top); the score at a uniform coverage across all samples (middle, 10 million reads for human and mouse, 1 million reads for Drosophila); the modeled quality score at infinite coverage (bottom, only samples with at least 4 million reads in human and mouse or 1 million reads in Drosophila). For each plot, the Pearson correlation coefficient and associated p-value are shown.

In addition to noise injection, we also examined the effects of heterogeneity on sample scores. Pairs of samples with quality scores differing by less than 1% at the 1 Mb resolution were mixed in varying ratios at 20 million read coverage. At all resolutions, samples showed decreased quality scores when part of a mixture, indicating that QuASAR is sensitive to heterogeneity as well as noise (Figure 2B). Thus, QuASAR quality scores not only detect noise but also the dilution of spatial consistency by superimposition of multiple disparate configurations.

Next, we examined the effects of coverage on quality scoring. As coverage decreased, quality scores decreased in a highly consistent manner for all samples and resolutions (Figure 2C). All quality score vs. log-transformed coverage relationships (for samples with at least 8 million reads for mouse and human samples, 1 million reads for Drosophila) were fit using logistic curves. This suggests that additional sequencing has diminishing returns on quality. Further, there exists some upper limit of quality for each sample. We also find that, consistent with expectations, the amount of coverage necessary to approach this quality asymptote increases with increasing resolution. In other words, a sample requires much less coverage to fully resolve large-scale conformational features in a consistent and high-quality manner. The X-axis offset of different samples also indicates that there should not be an arbitrary guideline for target coverage to resolve a particular resolution as the quality response to coverage is tissue and assay-specific. However, a minimum coverage based on currently tested samples may be appropriate.

Despite different resolutions plateauing at different rates, quality scores showed good agreement in sample ranking across resolutions (Figure S1A). Ranking was more consistent between more similar resolutions, indicating a greater overlap in the features being assessed compared to larger jumps in resolution. Because quality is a function of coverage, we also examined sample rankings at a uniform coverage level to remove any confounding effects. In most cases, the sample rankings still showed good agreement, although the consistency did decrease (Figure S1B). Interestingly, the mouse-derived samples showed an increase in correlation across resolutions after accounting for coverage.

We also examined how QuASAR quality scores related to various read statistics that have been used as proxies for sample quality. As observed in the coverage analysis, samples do not plateau at the same rate as a function of coverage. This means that a simple comparison across sample qualities at a given resolution may be misleading. In order to address this, we compared read statistics to three different quality reference points: the quality score at each sample’s actual coverage; the quality score at a uniform coverage level; and the inferred quality limit as determined by logistic curve fitting. Although we compared scores to four different read statistics, only the percentage of *cis*reads out of all valid reads showed a consistent and strong relationship to QuASAR quality scores (Figure 2D). For human and mouse samples, all three sets of quality scores showed strong correlation to the *cis*read ratio. However, the quality limit scores performed significantly better than the other two sets. The Drosophila samples did not show this same pattern and in fact negatively correlated to the *cis*read ratio. It is unclear why this pattern did not hold for Drosophila samples, although the number of samples is much lower compared to either mouse or human sample sets. The three other read statistics tested, percentage of reads with an insert size too large, percentage of reads from fragment circularization, and percentage of reads from putative missed restriction cuts showed no consistent relationship across any of the quality sets (Figure S2).

### Reproducibility results

To determine the performance of QuASAR replicate scoring, we began by finding replicate scores as a function of noise. In order to assess the effects of noise, we calculated a replicate score for each sample across varying resolutions and levels of noise injection into its matched biological replicate. The majority of samples showed a monotonically decreasing relationship between replicate score and noise level (Figure 3A). About 15% of sample-resolution relationships did not strictly hold to this trend. In all of these cases, the scores hovered around the noise-free replicate score before decreasing. Of these, most (35 of 42) occurred at the lowest level of resolution. None of these score fluctuations was over 1% above the noise-free replicate score. These results demonstrate the robustness of QuASAR replicate scores to noise, particularly when comparing macro features (low resolution).

**Figure 3:**
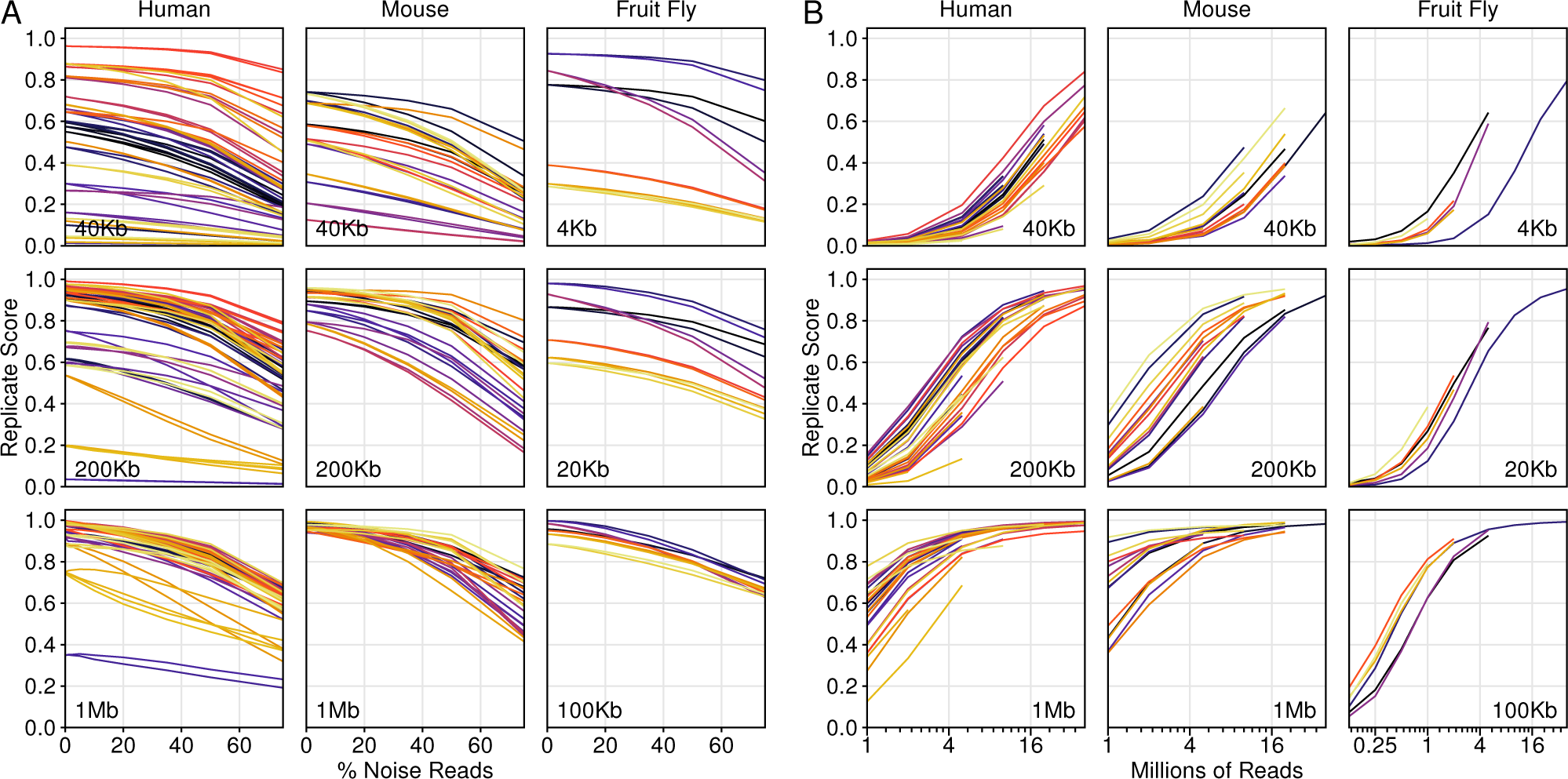
QuASAR replicate scores are sensitive to both noise and coverage. A) Replicate scores were calculated for each pair of sample replicates between the noise-free first replicate and the noise-injected levels of the second replicate and vice versa for each resolution. Sample color coding is consistent within each species across resolutions. B) For each pair of replicates, replicate scores were calculated at each coverage level and resolution. Sample color coding is consistent within each species across resolutions.

Next, we examined how coverage impacted replicate scoring. For each biological replicate sample pair, samples were down-sampled to various equal numbers of reads and replicate scores were calculated across multiple resolutions. In all sample pairs, replicate scores increased as a function of coverage following a logistic curve and ranging between zero and one (Figure 3B). Every sample pair showed a rightward shift of the curve midpoint as resolution increased, indicating that replicate agreement for macro features, such as compartments, occurred at lower coverage levels than fine-scale features, such as topologically associating domains or loops.

Finally, we calculated replicate scores for all pairwise combinations of samples within each species set across multiple resolutions. For human and mouse samples, pairs were scored at 1 Mb, 200 Kb, 40 Kb, and 10 Kb resolutions while Drosophila samples were scored at 100 Kb, 20 Kb, and 4 Kb (Figure S3). For all sample pairs, we took the highest score across all resolutions. The majority of biological replicate sample pairs scored close to one, the maximum replicate reproducibility score, indicating strong agreement in Hi-C signal between samples (Figure 4A-C). For all replicate pairs we also generated a pseudo-replicate sample composed of half of all reads from each replicate, randomly sampled and combined. Scores between samples and their pseudo-replicates were always higher than true biological replicates. In nearly all cases unrelated sample pair scores showed clear separation from replicate scores. There were two exceptions to this: samples with low biological replicate scores, and human ES and mesendoderm cells. The latter may be because the mesendoderm cells were differentiated from the ES cell line and either retain a strong conformational resemblance or differentiation was incomplete. We also observed elevated reproducibility scores for samples derived from the same tissue or cell line but from unrelated experiments. These samples typically fell in between the range of unrelated and biological replicate pair scores. Two exceptions to this pattern were human embryonic stem cells and mouse primary fetal liver cells, both of which showed reproducibility scores at or above biological replicate scores. We also observed that there was a relationship between sample quality scores and biological replicate scores such that if at least one of the replicates was of lower quality, the replication score was lower (Figure 4D).

**Figure 4:**
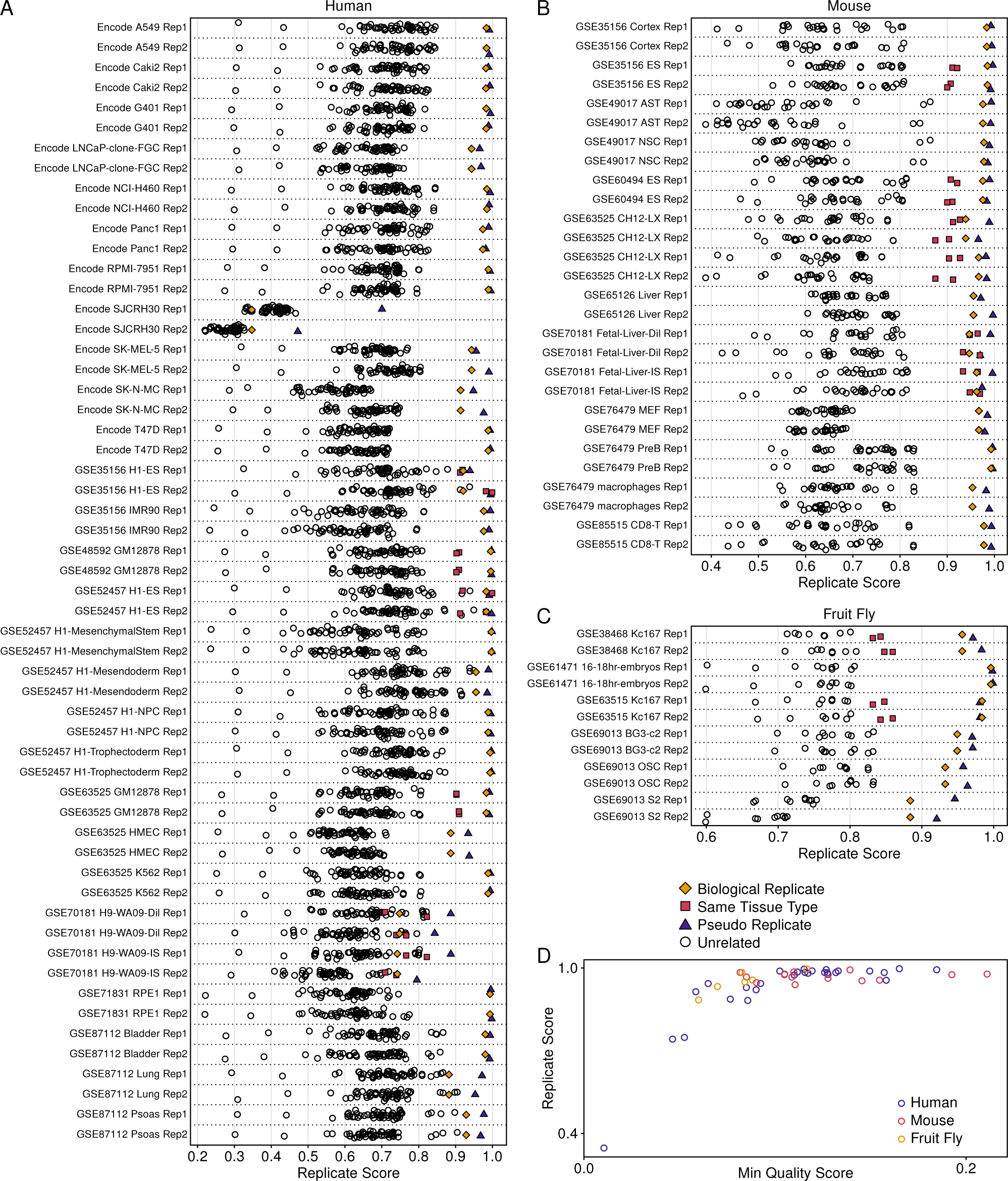
QuASAR replicate scores reflect both sample quality and consistency between sample origins. A-C) Replicate scores are denoted as points for each sample with an all to all pairwise comparison scheme within each set of species samples. Pairs include true replicates (gold diamonds), pseudo-replicates (purple triangles), same tissue of origin but non-replicates (fuchsia squares), and unrelated samples black circles). D) For each replicate pair, the lower of the two quality scores is plotted against the pair’s replicate score.

### Determining maximum resolution

One of the key choices to be made in using Hi-C data is determining an appropriate resolution for analysis. To answer this rigorously we propose a combination of quality and replicate scores to determine empirical cutoffs. In order to find the best cutoff values, we used an iterative process, cycling between using replicate or quality scores for classification followed by quality or replicate scores for cutoff value determination, respectively. For each each step, each sample and resolution combination tested was classified as passing or failing based on one set of scores (replicate or quality) and that set’s associated cutoff value. These labels were then applied to the other set of scores (quality or replicate) and a new cutoff value for that score type was determined based on minimizing the sum of the two Gini impurity indices for scores falling above and below the cutoff. This was repeated, reversing the score sets and cutoffs, until cutoff values stabilized. We tested initial replicate score cutoffs ranging from 0.75 to 0.99 and resolutions 10 Kb, 40 Kb, 200 Kb, and 1 Mb for mouse and human and 4 Kb, 20 Kb, and 100 Kb for Drosophila (Figure 5A). We found that across this range of initial replicate cutoffs, there were four sets of stable cutoff value pairs (Figure S4A). We selected the pair of cutoffs with the lowest combined Gini impurity score as our stringent cutoffs and the second lowest scoring pair as our loose cutoffs. For both cutoff sets, samples were partitioned into two groups with distinct distributions (Figures 5B & S4B) We then estimated resolution limits for each sample based on quality and replicate scores. For each score type, the resolution limit was defined as the point at which the log-transformed resolution vs. score curve equaled the score cutoff (Figure 5A). To further validate our cutoff values, we compared estimated resolution limits determined from quality and replicate analyses. Estimates from the two measures showed significant agreement for both cutoff sets (Figures 5C & S4C). The resolution limits also matched with a visual assessment of the data such that features were resolvable by eye at and above the resolution limits but were sparse or absent below (Figure 5D).

**Figure 5:**
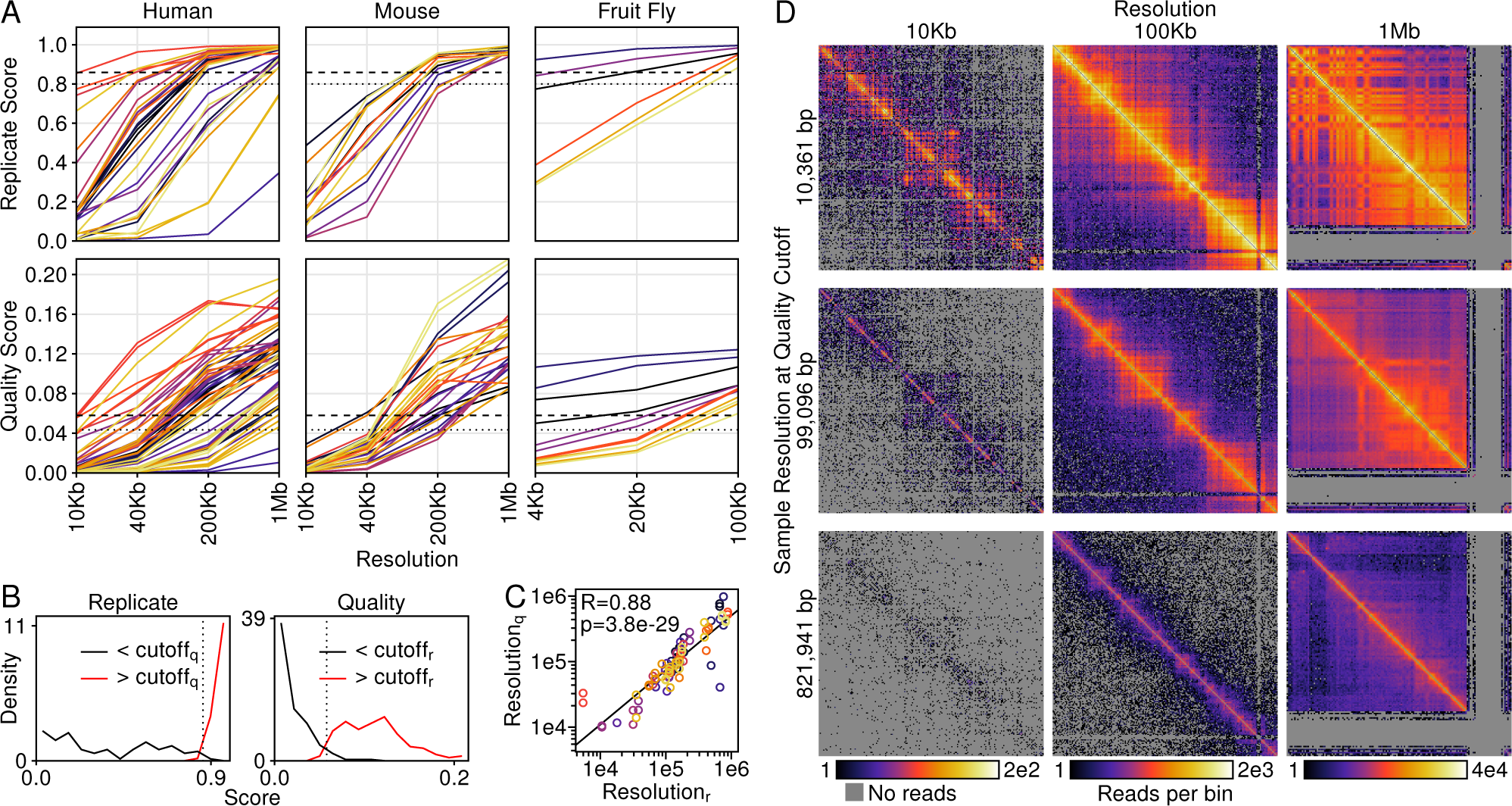
Calculating maximum usable resolutions from QuASAR scoring. A) QuASAR replicate (top) and quality (bottom) scores plotted as functions of log-transformed resolutions. Sample color-coding is consistent within each species plot pairs. Stringent and loose cutoff values are denoted by dashed line and dotted lines, respectively. B) Replicate (left) and quality (right) score distributions, partitioned by stringent quality and replicate cutoffs, respectively. Dotted lines donate the best separation point between partitions based on Gini impurity scoring. C) Resolution limits for each sample determined by stringent replicate and quality cutoffs. The Pearson R value and associated p-value are shown. D) Three samples (rows), selected for resolution limits close to target resolutions 10 Kb, 100 Kb, and 1 Mb, are shown binned at each target resolution (columns).

## Discussion

We have demonstrated the variability of Hi-C dataset quality across a variety of measures including resolution, coverage, and heterogeneity. Accurate quantification of this variability is necessary to make informed dataset selection and analysis choices. One of the primary strengths of QuASAR is its ability to provide resolution-specific information, which will benefit both producers and consumers of Hi-C data. It is currently difficult, if not impossible, to gauge the sequencing depth required to achieve a target level of resolution for analysis. QuASAR can provide crucial information for estimating the number of reads for a specific library necessary to produce high-confidence results at that target resolution by determining both the underlying library quality and the quality return on sequencing. For end users of Hi-C data, the redundancy of Hi-C datasets for specific cell lines or tissue types presents a challenge in selecting the most suited data for their analysis goals. An objective quality measure allows comparison of Hi-C protocols to evaluate strengths and weaknesses with respect to data quality. In addition to individual sample quality, the ability to find sample similarity between replicates or unrelated samples is of interest to the Hi-C community. Two recent publications describe tools to do just that [21, 22]. While both appear to function well, their outputs, while quantitative, give a binary-feeling classification of same/different. As we demonstrate in this study, there may be more nuance to sample comparisons (Figure 4). It is important not only to show the similarity of samples but to provide context such as at which resolutions the samples appear similar and are they similar enough to provide reliability to analyses. This is especially important given the prevalence of cell line-derived Hi-C data and the heterogeneity and instability of cell line genomes [23, 24]. In this study, we observed that Hi-C sample similarity was lower between cell line-derived datasets produced in different labs than between replicates, even at the lowest resolution of analysis. In many cases these scores approached the level of similarity seen for unrelated samples. This may serve as a cautionary tale about mixing dataset origins without verifying their similarity. Thus, to get the best return from time and financial investments in Hi-C data, it is important to evaluate the data critically prior to drawing conclusions. To this end, QuASAR provides an objective means of comparison of both individual samples and replicate agreement while providing context and limits to these measures.

## Methods

### Hi-C data processing and normalization

Hi-C raw read data were obtained from the Sequence Read Archive (SRA; https://www.ncbi.nlm.nih.gov/sra) or, in the case of ENCODE data, the ENCODE Data Portal (https://www.encodeproject.org; Table S1). Read ends were aligned using BWA mem version 0.7.12-r1039 and default settings [25] to the appropriate genome build (Table S1). Reads were considered valid and retained if both ends uniquely mapped to single locations, or at least one end spanned a ligation junction and mapped uniquely to two restriction fragments. In cases where both ends mapped to multiple fragments, reads were only kept of the upstream and/or downstream fragment locations matched across read ends. Reads were processed and normalized using HiFive version 1.3.2 [26]. Distance-dependent signal curves were estimated using the settings shown in Table S1. A maximum insert size of 650 bp was used to filter reads. Fends (fragment ends) were filtered to have a minimum of one valid interaction greater than 500 kb. Fend interaction filtering was applied only for the normalization step. All quality analyses were performed on unfiltered reads.

Data were normalized using the “binning” algorithm correcting for GC content, fragment length, and mappability. GC content was calculated from the 200 bp upstream of restriction sites or the length of the fragment, whichever was shorter. Mappability was determined using the GEM mappability function, version 1.315 [27]. Mappability of 36-mers was calculated every 10 bp with an approximation threshold of six, a maximum mismatch of 1 bp, and a minimum match of 28 bp. For each fend, the mean mappability score for the 200 bp upstream of the restriction site, or total fragment size if smaller, was used. For normalization, only intra-chromosomal reads with an interaction distance of at least 500 kb were used. GC content and fragment length were partitioned into 20 bins each and mappability was partitioned into 10 bins. All parameter partitions were done such that together they spanned the full range of values and contained equal numbers of fends in each bin. All bins were seeded from raw count means and GC and length parameters were optimized for up to 100 iterations or until the change in log-probability was less than one, whichever was achieved first. Normalization was used only for noise model construction.

### QuASAR matrix transformation and scoring

All intra-chromosomal raw reads were binned at a predefined resolution (1 Mb, 200 Kb, 40 Kb, or 10 Kb for mouse and human datasets, 100 Kb, 20 Kb, or 4 Kb for Drosophila datasets), depending on the analysis. Only numbered chromosomes and the X chromosome were used for analysis in human and mouse datasets while chromosomes 2L, 2R, 3L, 3R, 4, and X were used for fly datasets. For each chromosome, only rows and columns that had at least one read for an interaction occurring over a span of 100 bins or fewer were included. All other rows and columns were marked as invalid. The resulting matrix is defined as *R*. A scaled matrix, *S*, was calculated as the square-root of the sum of matrix *R* plus one. A distance-normalized matrix, *N*, was calculated by dividing each diagonal of *R* by the sum of the diagonal divided by the sum of valid rows/columns in that diagonal. A local correlation matrix, *C* was then calculated from *R* such that for each pairwise set of rows no more than 100 rows apart, the correlation was calculated between valid column entries not more than 100 columns from either row number, not including self-interactions or the interaction between the pair of bins being correlated:

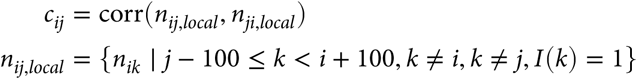

where *I*(*k*) is an indicator function taking on the value of one for valid rows/columns, and zero otherwise. Entries in matrix *C* were considered valid if there were at least three points in the correlation and the standard deviation of both *n*_*i,local*_ and *n*_*j,local*_ were non-zero. The QuASAR transformed matrix, *T* was the element-wise product of *S* and *C*.

QuASAR quality scores were calculated as the sum of valid elements of the transformed matrix *T* divided by the sum of the corresponding elements of *S* minus the mean of the corresponding elements of *C*. Thus, with the indicator function *I* (*i*, *j*) taking a value of one for valid element *c_ij_* and zero for an invalid element, the Quasar quality score is defined as follows:

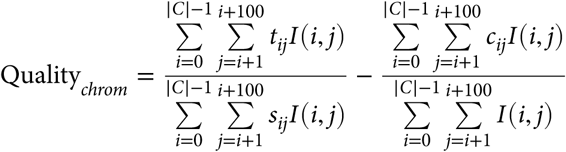

This score was calculated on a per chromosome basis. A global quality score was calculated by finding each of these sums across across all chromosomes prior to division.

QuASAR replicate scores were calculated by finding the correlation of the two sample transformation matrices *T* for all elements that are valid in both matrices. These scores were calculated on a chromosome by chromosome basis as well as a global score across all valid matrix elements for all chromosomes.

### Hi-C noise and low-coverage modeling

The noise model employed was based on all noise coming from random ligation. For each genome and restriction enzyme combination, the median bin correction values across all relevant datasets were used to calculate expected values for all bins at the lowest resolution used for analysis (10 Kb for human and mouse data, 4 Kb for Drosophila data). Bins that had zero observed reads in any of the used datasets were set to zero. Expected values were converted into probabilities by dividing values by the sum of all expected values. For noise-injected sample, a random fraction of reads corresponding to the target noise percentage were randomly selected and discarded. The same number of reads were then drawn from the above described noise distribution and combined with the remaining sample reads. This was done prior to filtering or normalization.

Low coverage samples were generated by random selection and removal of reads prior to any filtering or normalization.

### Pseudo-replicate and heterogeneous sample generation

Pseudo-replicates were generated by random selection of reads from each replicate across all chromosomes prior to filtering or normalization. For each replicate, half of the reads were selected and combined, meaning that pseudo-replicates had a number of reads equal to the mean of the two replicates. Heterogeneous samples were produced the same way, although the percentage of reads drawn from each sample was varied from 0 to 100% at 20% steps.

### Coverage curve estimation

QuASAR quality curves as a function of coverage were calculated using the curve_fit function from the python package SciPy [28]. All lines were estimated using the logistic function as follows:

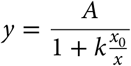

where *A* is the quality score upper bound, *x*_0_ is the inflection point coverage level, and *k* is a scale factor. Initial values for each sample were set to twice its maximum quality score, its maximum coverage, and 0.5 for *A*, *x*_0_, and *k*, respectively. *A* was bounded between -2 and 2, while the other parameters had no limits.

### Resolution cutoff calculation

Resolution cutoffs were calculated in an iterative fashion. Initial resolution cutoff values ranged from 0.750 to 0.990 at steps of 0.005. Each sample/resolution combination was labeled passing if its associated replicate score was greater than the initial resolution cutoff. Quality scores were then ordered and potential quality cutoff points were defined as the midpoints between adjacent quality scores. For each potential quality cutoff, the sum of the Gini impurity scores for the two partitions of samples (above and below quality cutoff) was calculated based on replicate cutoff labels, weighted by the number of samples in each partition. This quality cutoff was then used to partition associated replicate/resolution pairs. For all replicates, the lower quality score was used for label determination. The same procedure was followed for finding the new replicate cutoff value as described for the quality cutoff value. This process was repeated until both replicate and quality cutoff values remained constant. Maximum resolution limits were calculated based on quality or replicate curves as a function of the log-transformed resolution. For each sample or replicate pair, the resolution associated with the point at which the quality or replicate cutoff value, respectively, fell on the curve was used as the maximum resolution limit.

## Availability of data and materials

The datasets analyzed during the current study are available at https://bx.bio.jhu.edu/data/quasar. Analysis code and results are available at https://github.com/bxlab/Quasar_PaperAnalysis. QuASAR is packaged as part of the HiFive suite of tools (https://github.com/bxlab/hifive).

## Acknowledgements

Thanks to all members of the Taylor lab for useful discussion and feedback. Thanks to all of the members of the ENCODE 3D Nucleome subgroup for discussions on reproducibility and quality analyses. MEGS and JT were funded in part by NIH/NIDDK grant R24 DK106766 and NIH/NHGRI grant U41 HG006620.

## Supplementary Material

**Figure S1:**
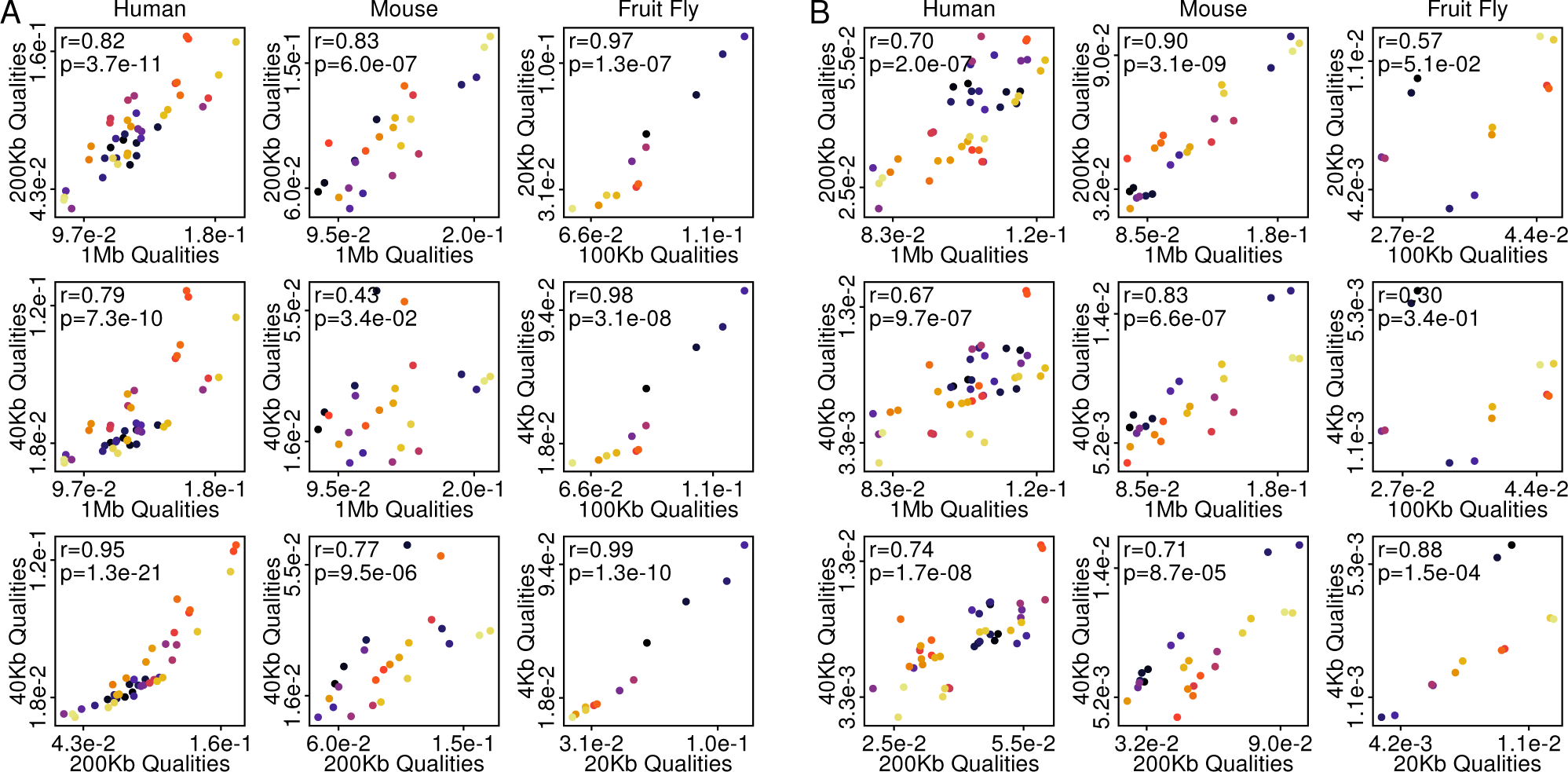
QuASAR quality scores show decreased consistency between larger changes in resolution. A) QuASAR quality scores from unaltered datasets are plotted between different pairs of analysis resolution. For each plot, the Spearman rank-order coefficient of correlation and associated p-value are shown. Sample color coding is consistent within a species across plots. B) QuASAR quality scores from samples down-sampled to a uniform coverage level are shown for different pairs of analysis resolution. For human and mouse data, all samples contain 10 million *cis*reads while for Drosophila samples each contain 1 million *cis*reads. For each plot, the Spearman rank-order coefficient of correlation and associated p-value are shown. Sample color coding is consistent within a species across plots.

**Figure S2:**
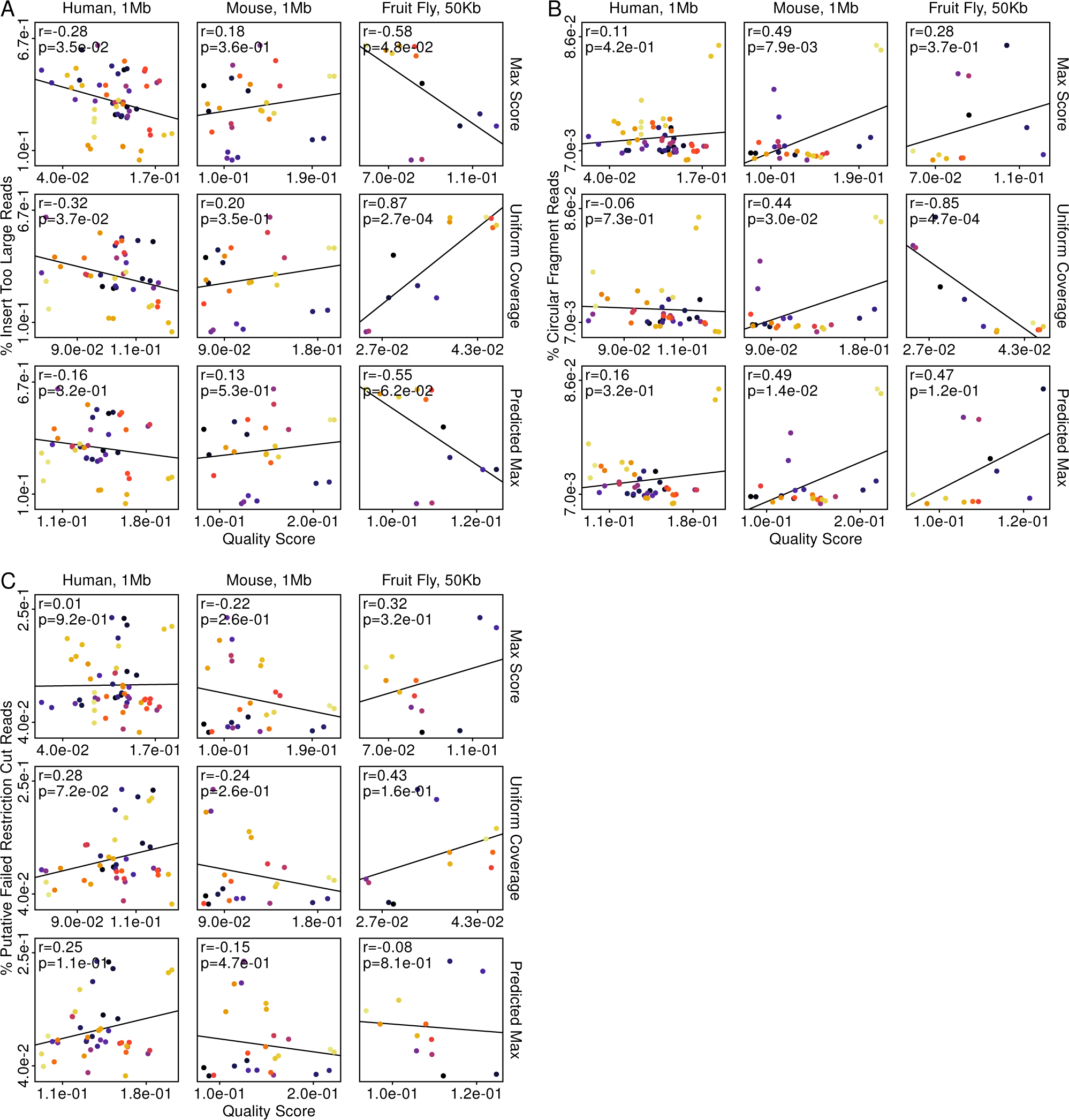
QuASAR quality scores show no consistent relationship to many common descriptive Hi-C statistics. QuASAR quality scores versus common Hi-C mapping statistics. For each panel, quality scores were derived from unaltered samples (top, 1 Mb resolution for human and mouse, 100 Kb resolution for Drosophila), samples down-sampled to equal coverage (middle, 10 million *cis*reads for human and mouse, 1 million *cis*reads for Drosophila), and modeled scores under infinite coverage (bottom, samples with at least 4 million *cis*reads for human and mouse or 1 million reads for Drosophila). For each plot, the Pearson correlation coefficient and associated p-value are shown. A) Quality scores are plotted versus the percentage of reads in each sample with an estimated insert size larger than 650 bp. B) Quality scores are plotted versus the percentage of reads circularized, or self-ligated restriction fragments for each sample. C) Quality scores are plotted versus the percentage of reads for each sample on adjacent restriction fragments that whose ends were in opposing orientations, allowing for the possibility of a failed restriction cut and circularization.

**Figure S3:**
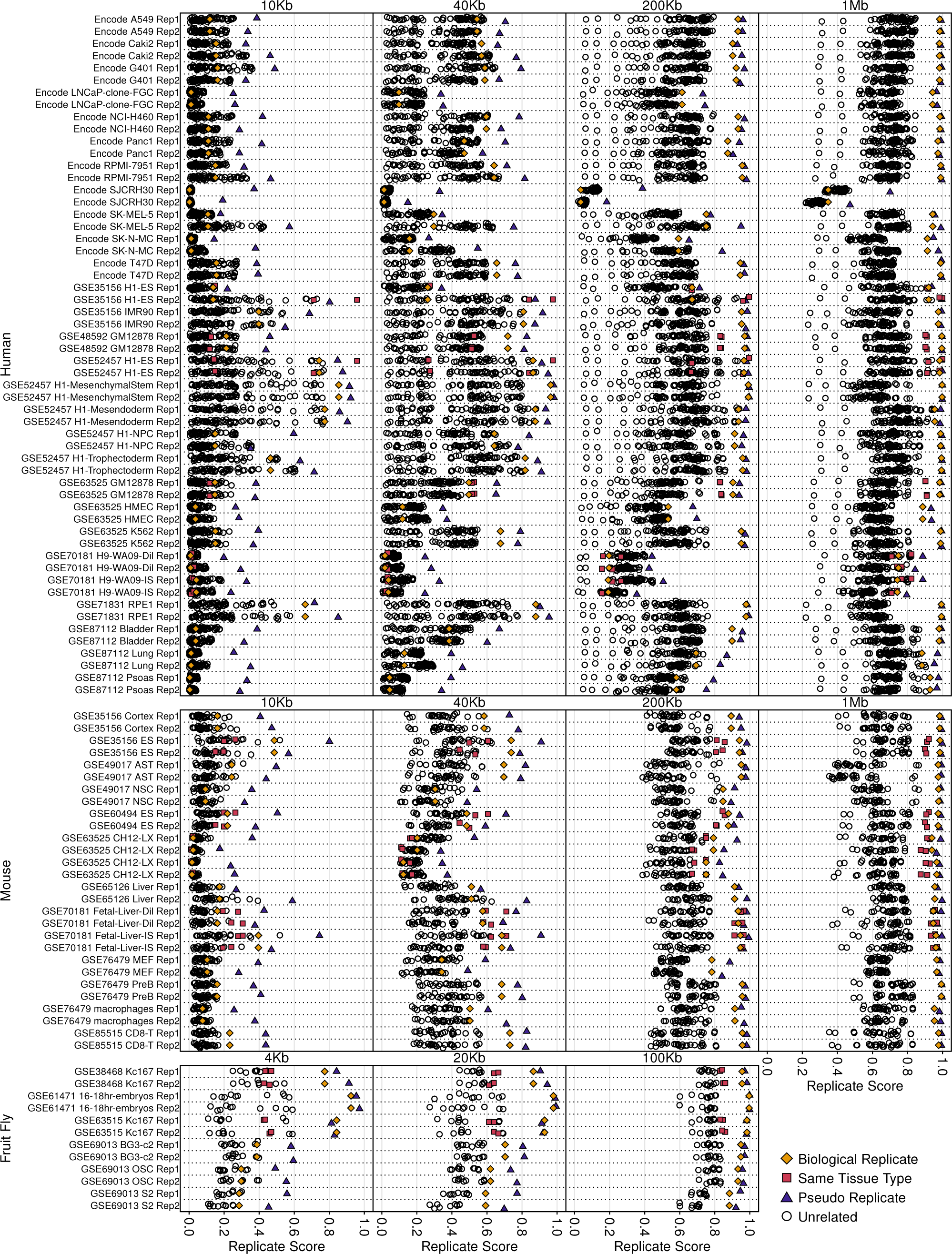
QuASAR replicate scores across all tested resolutions. A-C) Replicate scores are denoted as points for each sample with an all to all pairwise comparison scheme within each set of species samples. Pairs include true replicates (gold diamonds), pseudo-replicates (purple triangles), same tissue of origin but non-replicates (fuchsia squares), and unrelated samples black circles). For each species, the associated resolution is displayed at the top of each block of samples.

**Figure S4:**
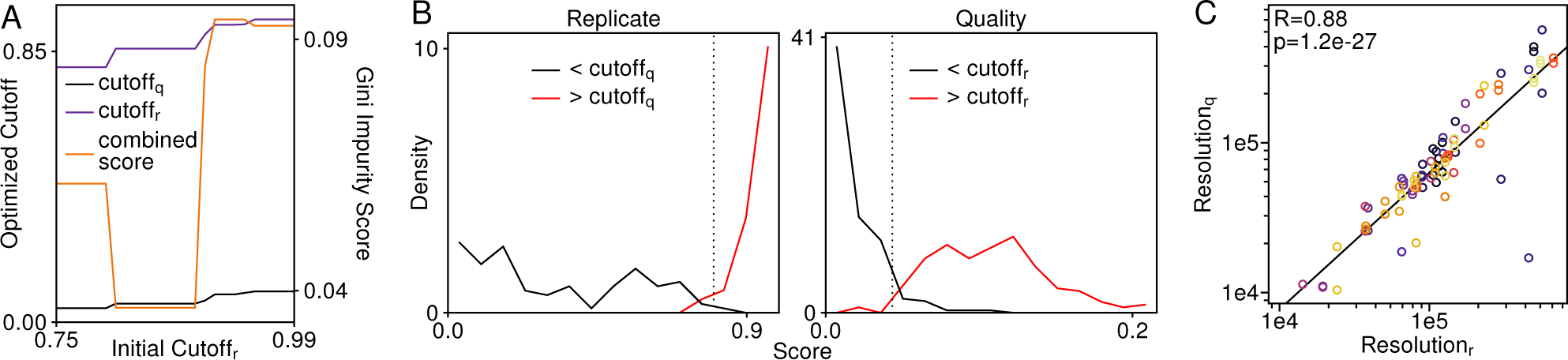
Determining and validating QuASAR score cutoffs for finding maximum usable resolutions. A) Optimized cutoff values and associated Gini impurity score sums as a function of starting replicate cutoff. B) Replicate (left) and quality (right) score distributions, partitioned by loose quality and replicate cutoffs, respectively. Dotted lines donate the best separation point between partitions based on Gini impurity scoring. C) Resolution limits for each sample determined by loose replicate and quality cutoffs. The Pearson R value and associated p-value are shown.

**Table S1:**
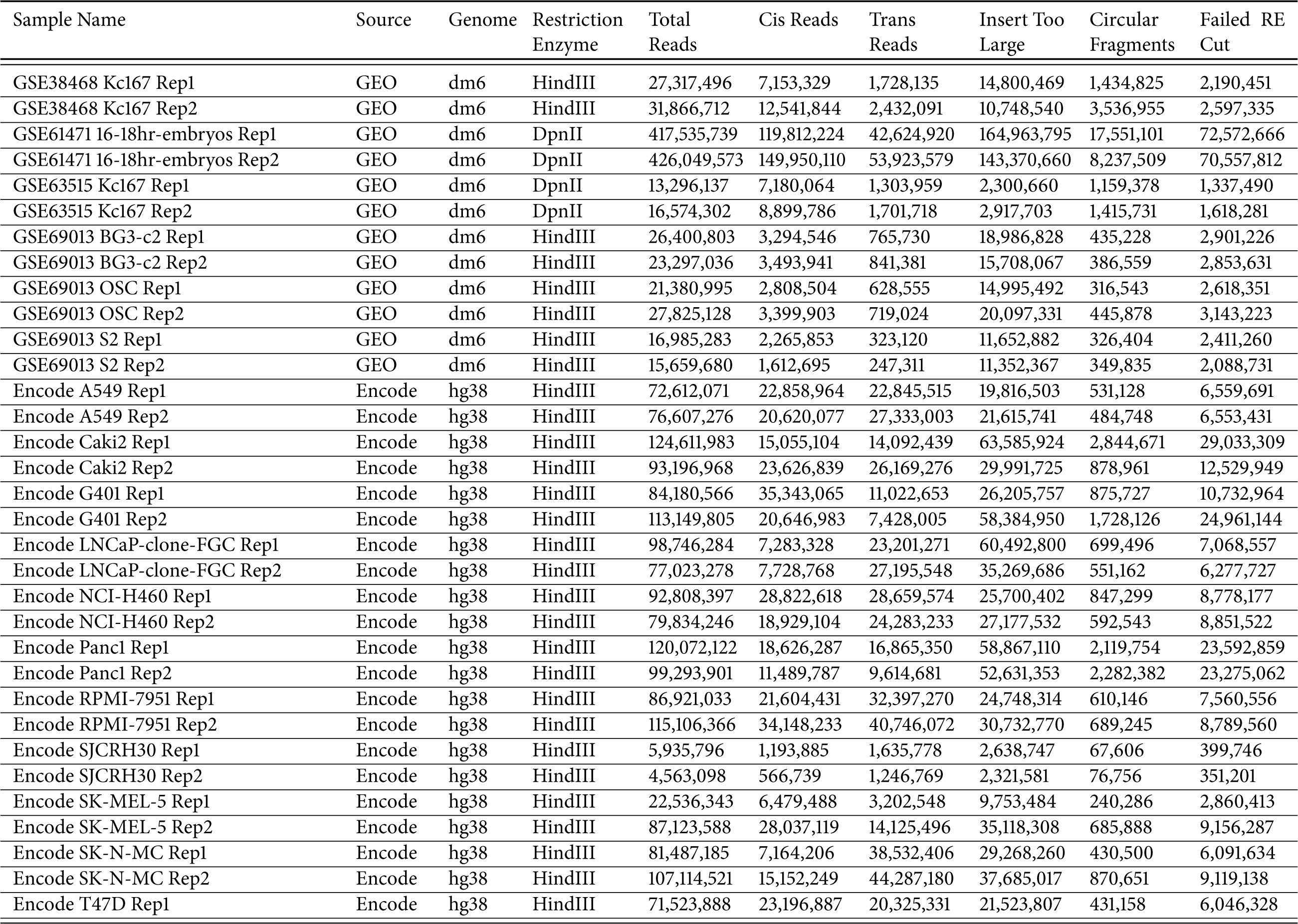
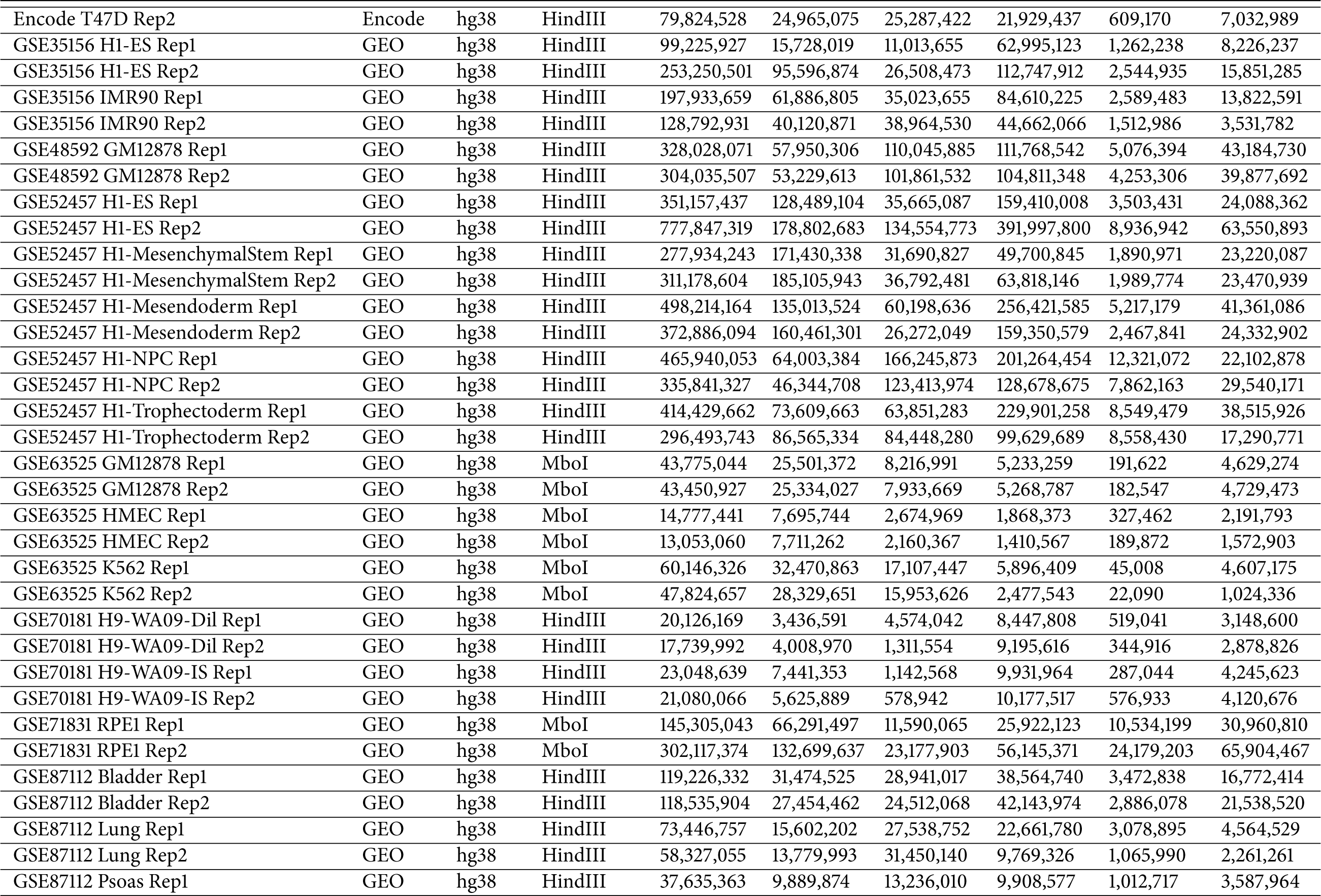
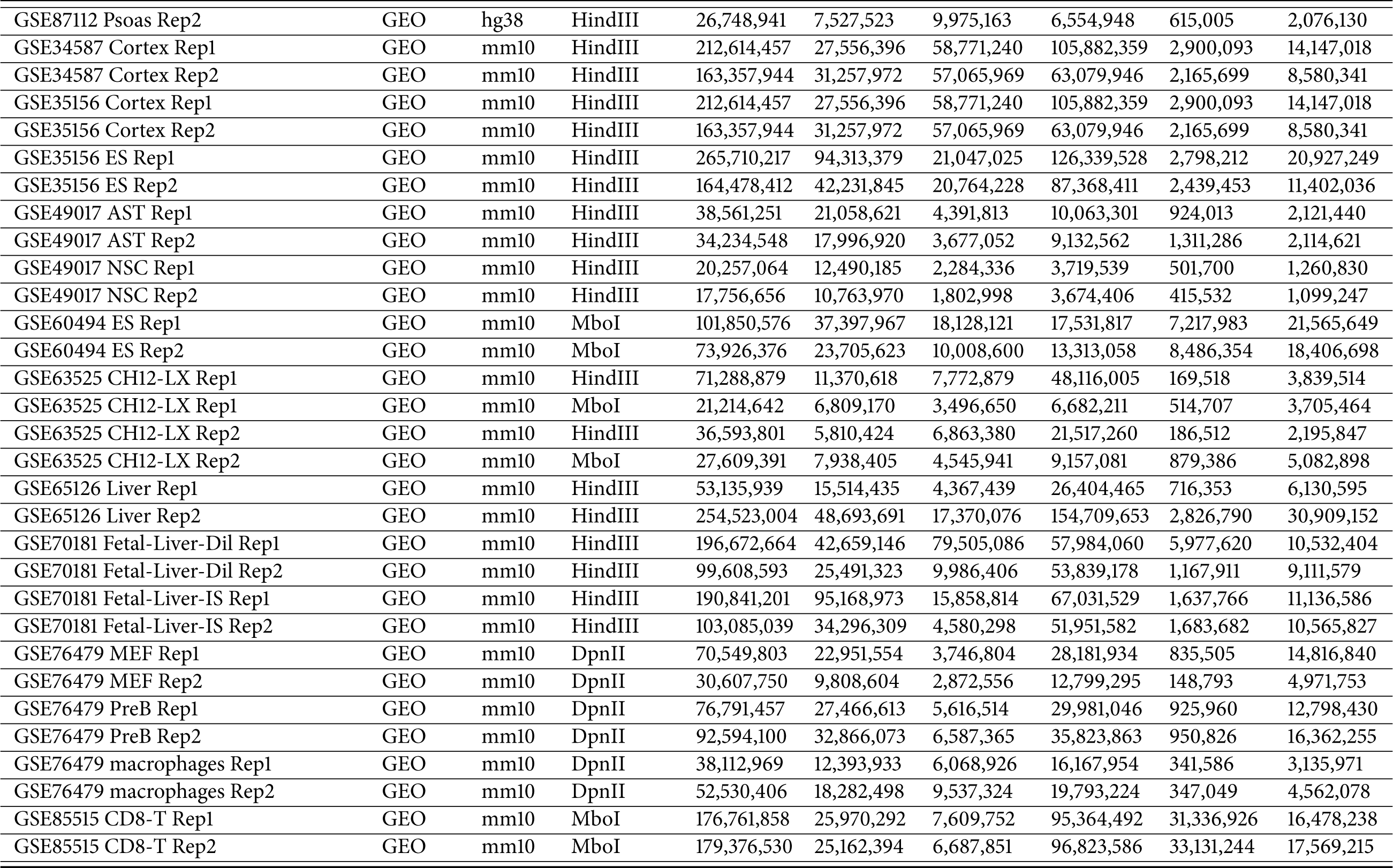
Datasets and associated read statistics used for analysis.

